## Supplemental figure and table for "Specific PIP_2_ Binding Promotes Calcium Activation of TMEM16A Chloride Channels"

[2]Department of Biochemistry and Molecular Biology

University of Massachusetts

Amherst, MA 01003, USA

### Supplementary Tables

**Table S1. Summary of atomistic simulations of TMEM16A**

| Systems | $Ca^{2+}$ | $PIP_2$ | Initial Structure | Length (μs) |
|---|---|---|---|---|
| *sim 1-3* | + | + | 5oyb | sim 1: 3.0<br>sim 2,3: 1.5 |
| *sim 4-6* | + | - | 5oyb | sim 4: 3.0<br>sim 5,6: 1.5 |
| *sim 7-9* | - | + | 5oyg | 1.0 each |
| *Meta 1-4** | + | + | *sim1*, 0.65 μs | 0.2 Each |

* metadynamics simulations in the $Ca^{2+}$/$PIP_2$-bound open state of TMEM16A.

### Supplementary Movies

**Movie S1**: Spontaneous opening of the $Ca^{2+}$-bound TMEM16A pore induced by specific $PIP_2$ binding. The movie was generated based on *sim1*. Only *chain B* is shown in cartoons, with TMs 3-4 colored in blue, TMs 5-6 in red and the rest in light green. The phosphate atoms in lipid head groups are represented as transparent spheres, and water molecules near the pore shown in sticks. The bound $PIP_2$ molecule is shown in ball-and-stick. The two bound $Ca^{2+}$ ions are shown as two red spheres. The inner gate residues, L547, S592 and I641, are shown as yellow sticks.\

**Movie S2**: The $Ca^{2+}$-bound TMEM16A pore remained collapsed without $PIP_2$. The movie was generated based in *Chain B* from *sim4*. See the Movie S1 caption for details of molecular rendering.

**Movie S3**: The $Ca^{2+}$-free TMEM16A pore remained collapsed even with specific binding of $PIP_2$. The movie was generated based in *Chain A* from *sim7*. See the Movie S1 caption for details of molecular rendering.

**Movie S4**: Spontaneous $Cl^-$ permeation through the open pore of TMEM16A. The movie was generated based on *sim3*. The permeating chloride is shown as a red sphere with coordinating waters shown in sticks. See the Movie S1 caption for details of molecular rendering.

### Supplementary Figures

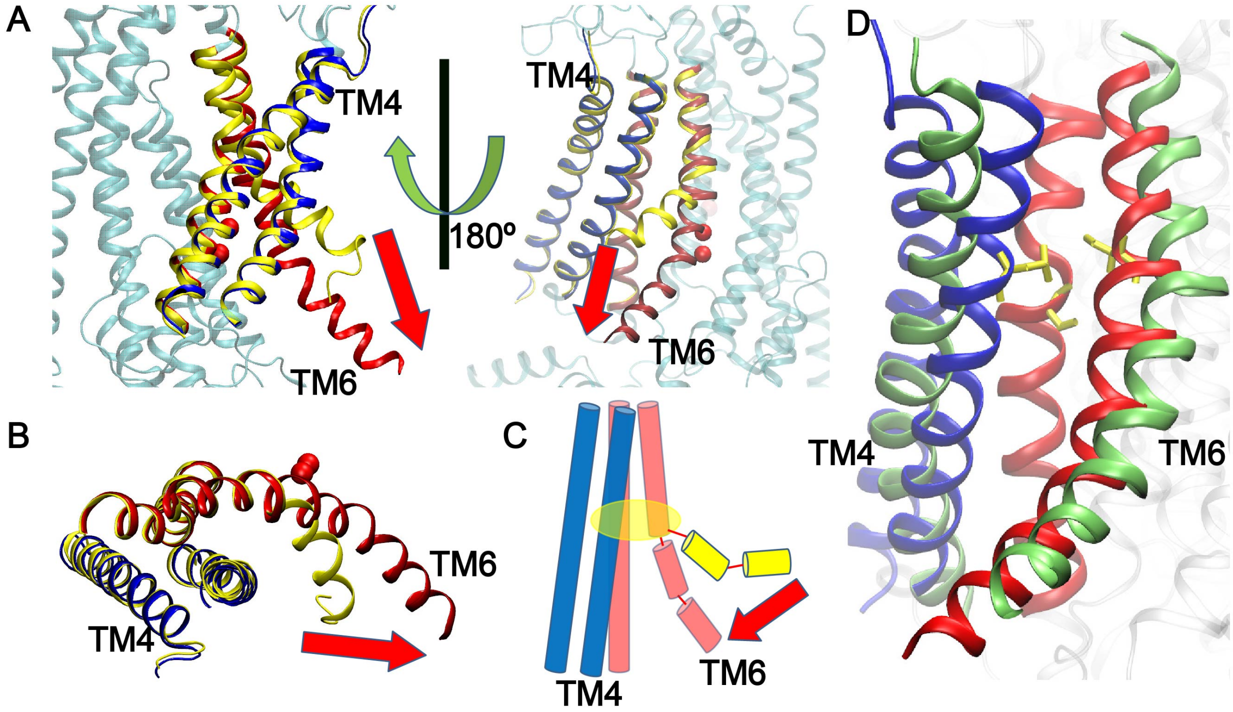


**Fig. S1.** $Ca^{2+}$-induced lower pore opening in TMEM16A. **A** and **B**, side and top views of the TMEM16A in the $Ca^{2+}$-free (PDB: 5oyg) and bound (PDB: 5oyb) states. TMs 3-4 and TMs 5-6 in the $Ca^{2+}$-bound structure are colored in blue and red, respectively; TMs 3-6 in the $Ca^{2+}$-free structure are colored in yellow. The movement of TM6 upon $Ca^{2+}$ binding is highlighted with the red arrow. **C**: cartoon illustration of the movement of TM6 upon $Ca^{2+}$ binding. **D**: Superimposed structurs of $Ca^{2+}$-bound TMEM16A with the fully opened nhTMEM16 lipid scamblase (PDB: 4wis). TM4 and TM6 of nhTMEM16 (green cartoons) are further separated than those of TMEM16A.

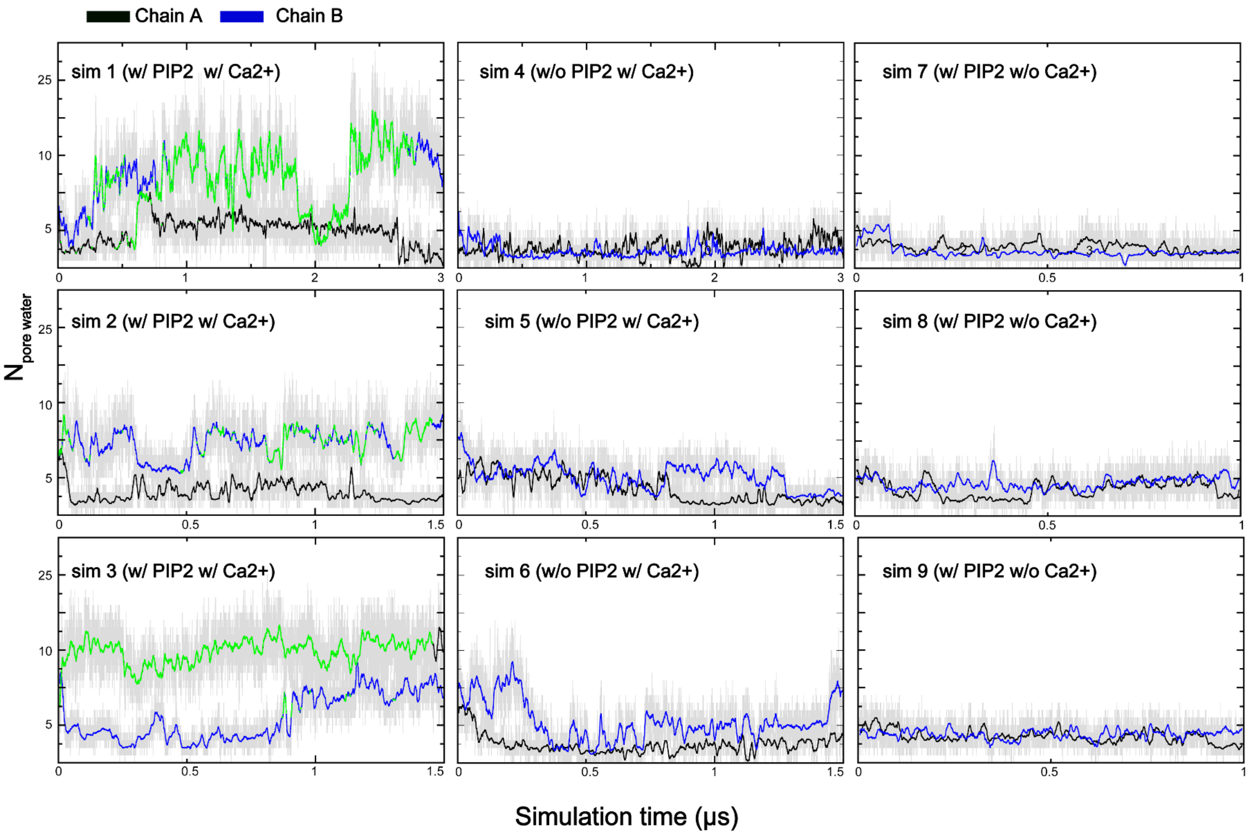


**Fig. S2** Number of pore water molecules in the neck region as a function of the simulation time for $Ca^{2+}$-bound TMEM16A with $PIP_2$ (*sim1-3*), $Ca^{2+}$-bound TMEM16A without $PIP_2$ (*sim4-6*), and $Ca^{2+}$-free TMEM16A with PIP2 (*sim7-9*). All snapshots that belong to the opened state cluster (see Fig. S3) are highlighted in green. Note that TMEM16A remained in deactivated states and never sampled any open state conformation throughout all simulations without both $Ca^{2+}$ and $PIP_2$ bound (*sim4-9*).

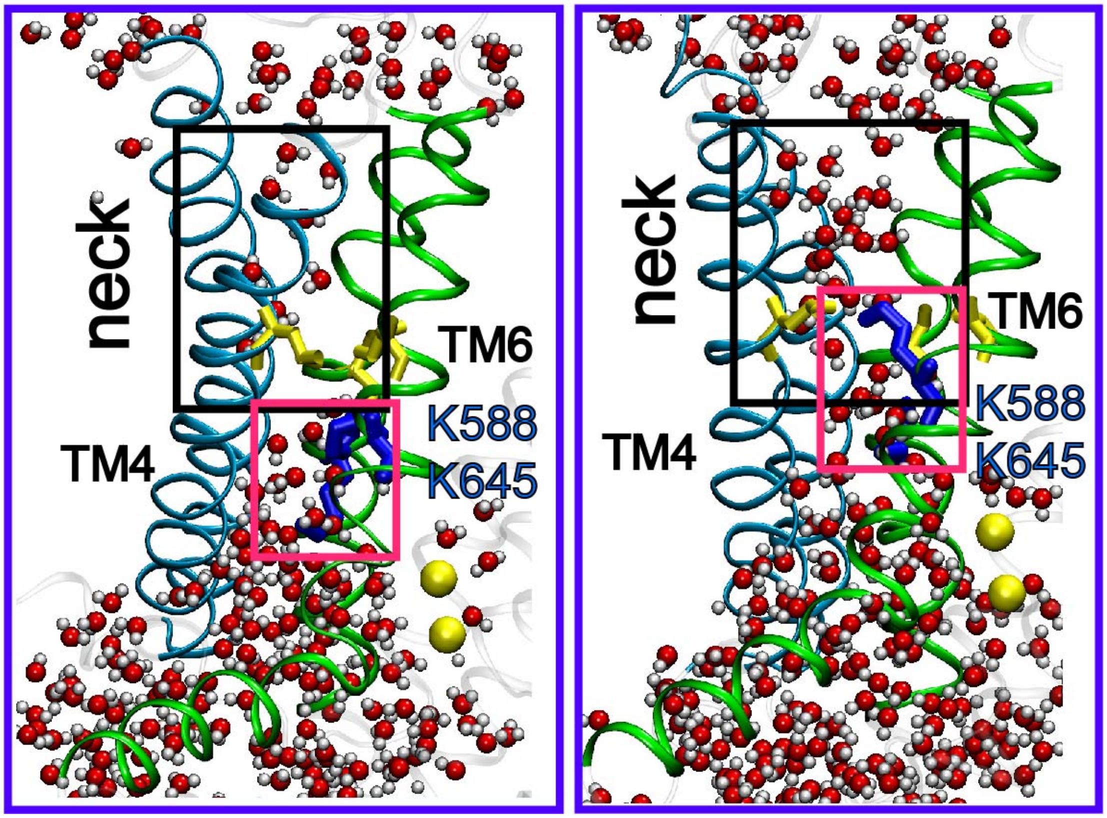


**Fig. S3** Representative snapshots showing pore hydration in the collapsed state (left, $Ca^{2+}$-bound but without $PIP_2$; *sim4,* 0.535 μs) and the dilated state (right, $Ca^{2+}$ and $PIP_2$ bound; *sim1,* 2.681 μs). TMs 3-4 and 5-6 are represented as cyan and green cartoons, respectively. Bound calcium ions are shown as yellow spheres and water molecule inside the pore as spheres colored by atom types (red: oxygen; white: hydrogen). The inner gate residues, L547, S592 and I641, are shown as yellow sticks. Two conserved basic residues below the gate, K588 and K645, are represented as blue sticks and highlighted with red boxes. The neck region is highlighted with a black box.

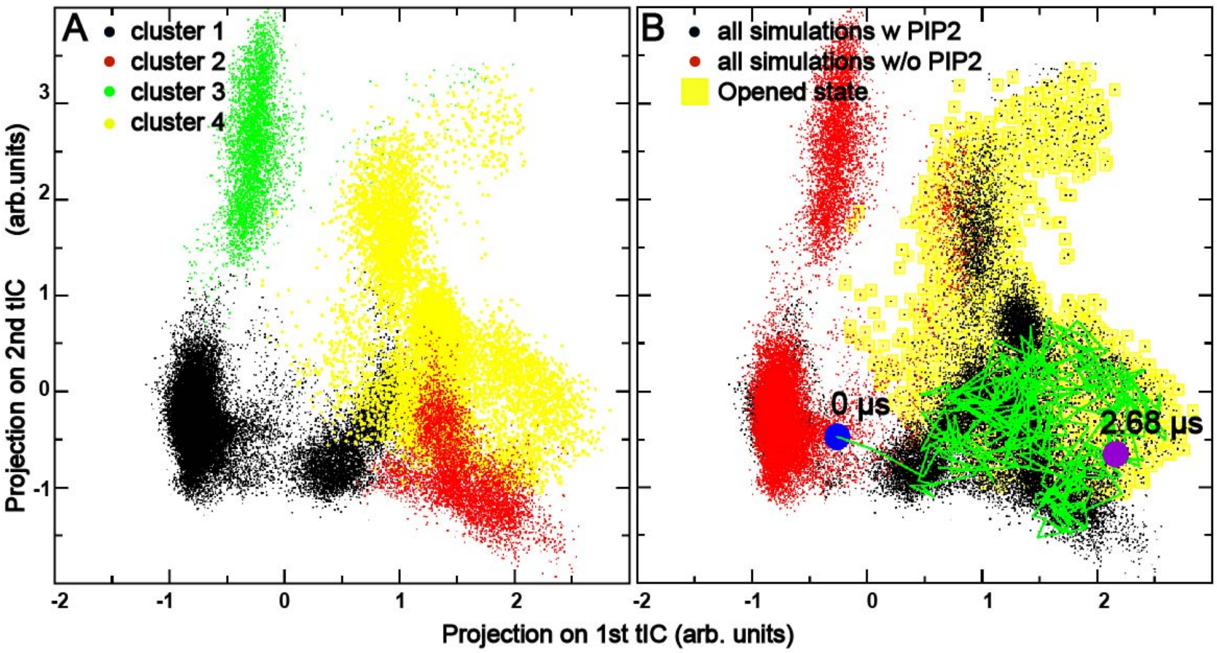


**Fig. S4** Clustering of TMEM16A pore conformational space. All snapshots sampled in trajectories s*im1-6* ($Ca^{2+}$-bound state with and with $PIP_2$) were clustered based on time-lagged independent component analysis (tICA) (see Methods for details). The snapshots are projected onto the first two tICAs. **A**: The clusters are colored by cluster IDs (1-4). The cluster 4 (yellow) is the open and conductive state. **B**: The clusters are colored by simulation conditions, with snapshots from simulations of $Ca^{2+}$-bound TMEM16A with and without PIP2 colored in red and black, respectively. The conformational space covered by the open state cluster (cluster 4 in panel A) is marked using a yellow background. The location of the initial structure (PDB: 5oyb) is labeled using the blue circle, and the location of the selected open state pore structure (around 2.68 $\mu s$ from *sim 1,* Chain B) is labeled using the purple circle. The green trace plots the trajectory of the pore structure of chain B sampled during *sim1*.

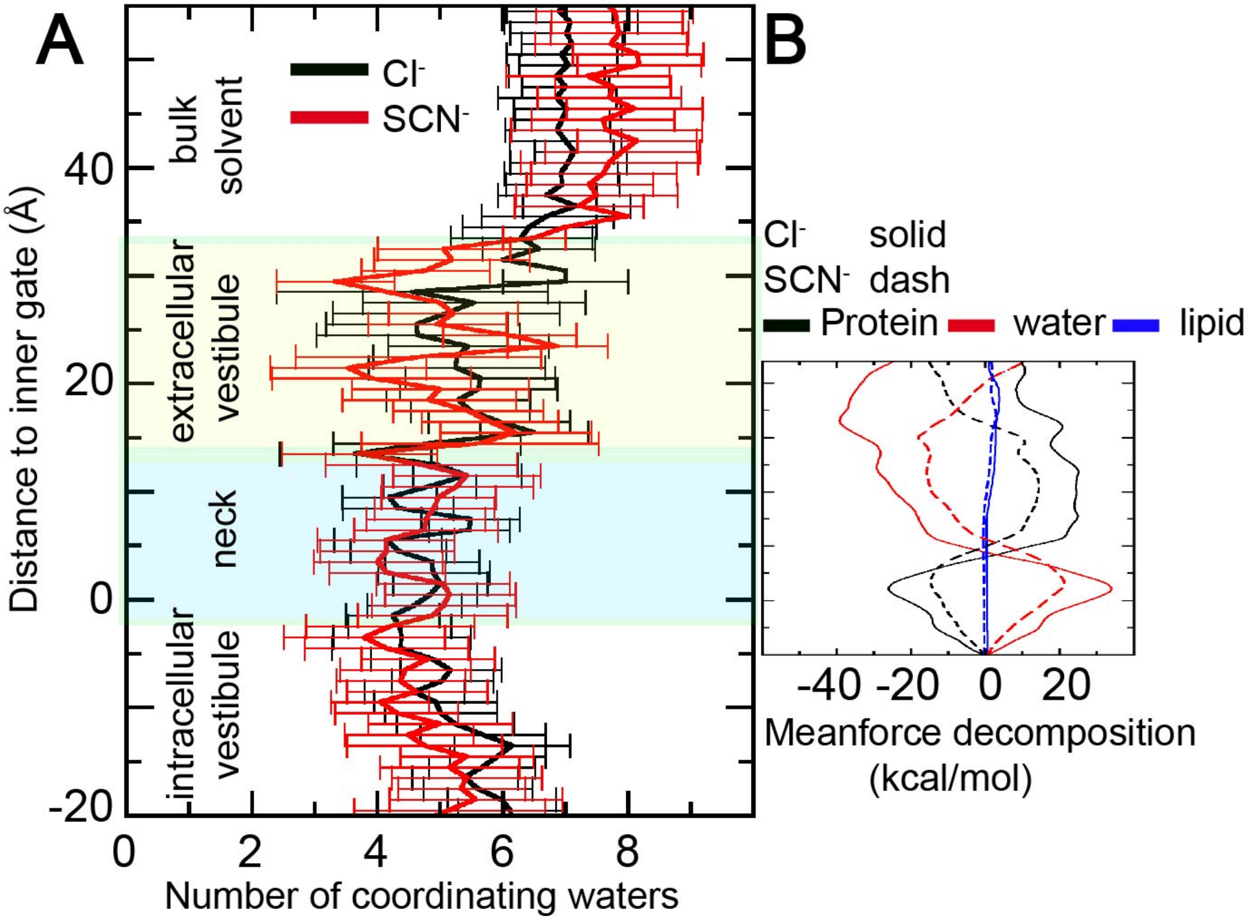


**Fig. S5** Solvation property and free energy decomposition of $Cl^-$ and $SCN^-$ permeation. **A**. The number of solvation waters of anions inside the channel, calculated as the number of water hydrogen atoms within 2.8 Å of $Cl^-$ (black trace) or $SCN^-$ (red traces). The results were calculated as the average from umbrella sampling trajectories and the error bars shown are the standard deviations. **B.** Contributions of protein, water and membrane to the free energy of anion permeation calculated using mean force decomposition. Various decomposed contributions are shown in solid and dashed lines for $Cl^-$ and $SCN^-$, respectively.

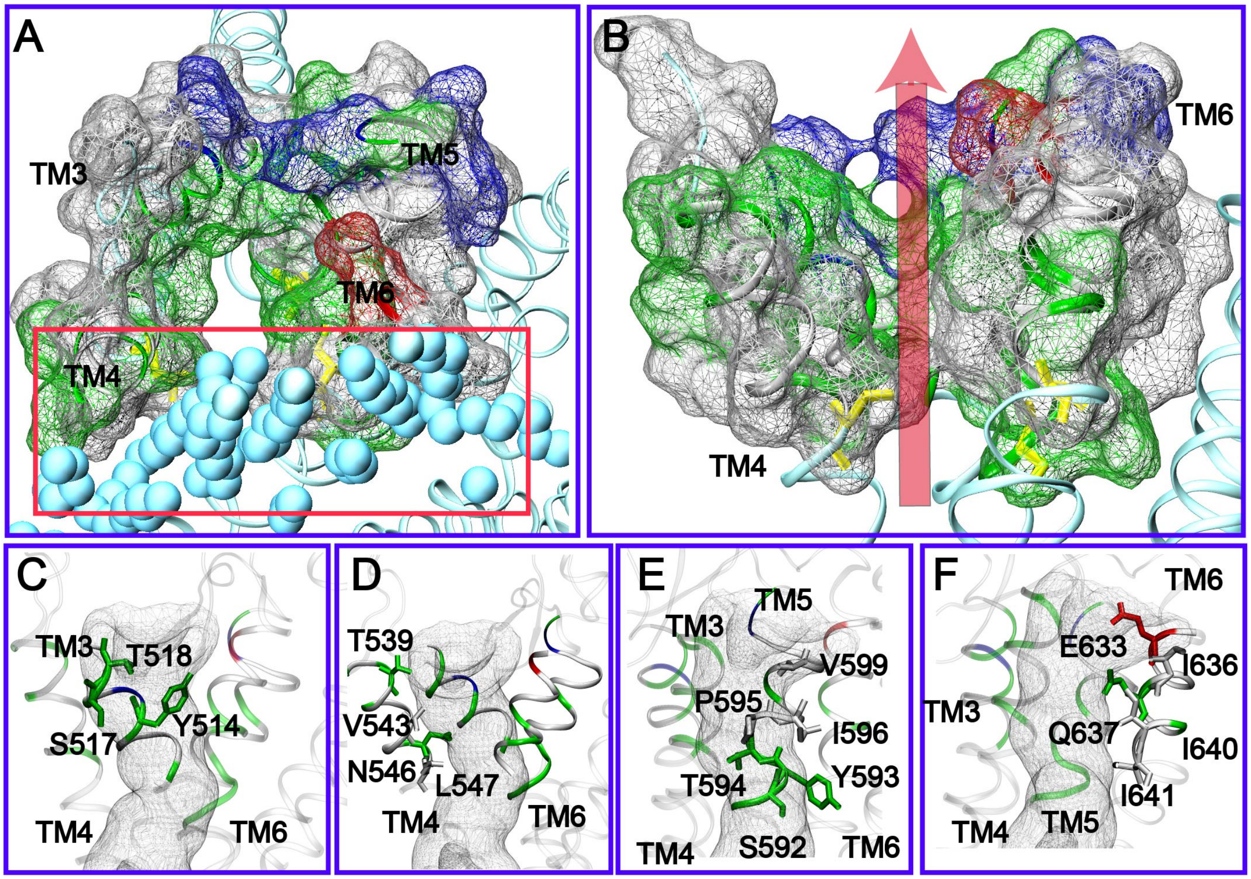


**Fig. S6.** Structural features of the open state of TMEM16A conduction pore. **A, B**: top and front side views of the pore surface. The van der Waals surface of the protein is represented as mesh surfaces. The hydrophobic, hydrophilic, positively charged and natively charged residues are colored in white, green, blue and red, respectively. The inner gate residues, L547, S592, I641, as yellow sticks. POPC lipid tails near TMs 4 and 6 (<9 Å heavy atom distance) are shown as cyan spheres in panels A and B. **C**-**F**: pore-lining residues on TMs 3-6, respectively. The snapshot is taken from *sim1* at 2.681 μs. The residues are colored by residue type (hydrophobic: white; hydrophilic or polar: green; negatively charged: blue; positively charged: red). The profile of the conducting pore calculated using HOLE is shown as the transparent tunnel in panels **C**-**F**.

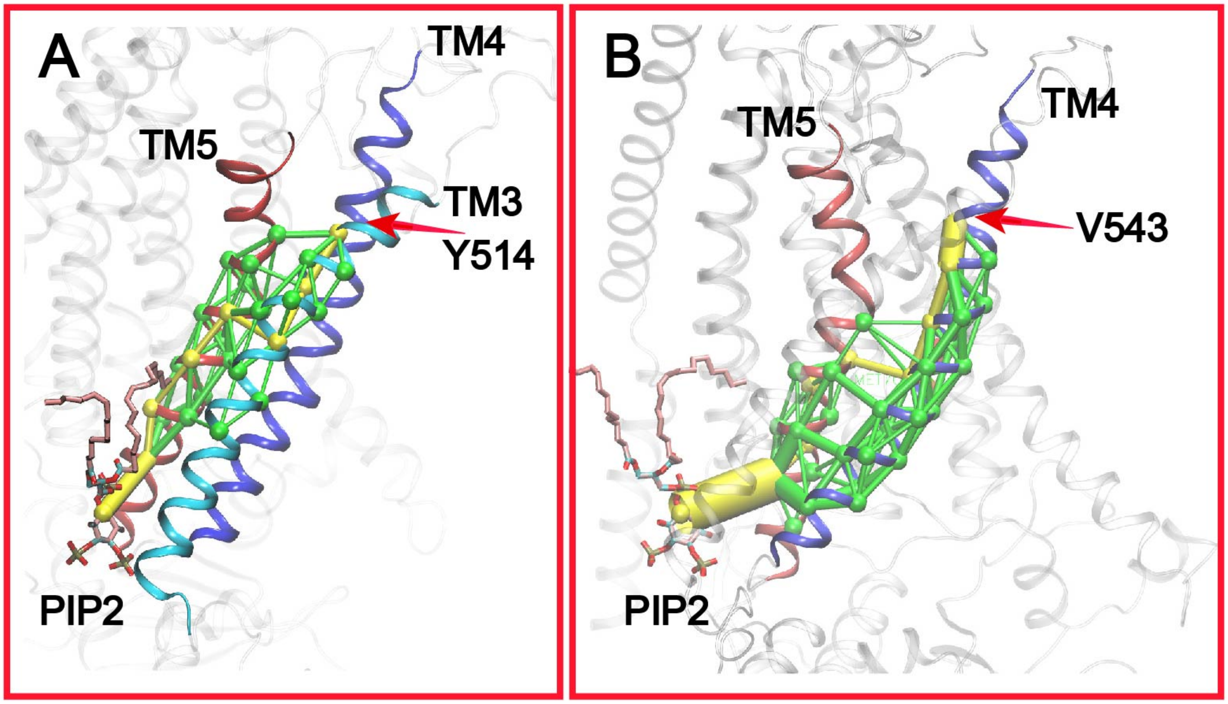


**Fig. S7** Optimal and suboptimal pathways of coupling between PIP2 and the neck region of the TMEM16A pore. **A**: Coupling pathways between PIP2 and Y514 in TM3, and **B**: between PIP2 and V543 in TM4. TMs 3, 4 and 5 are represented as cyan, blue and red cartoons, respectively. TM3 is not colored in panel B for clarity. The residues involved in the pathway are presented as green spheres and the contacts are presented as sticks. The optimal and suboptimal pathways are colored in yellow and green, respectively. The thickness of the edges represents the number of paths crossing that edge. See Methods for details of coupling pathway analysis.

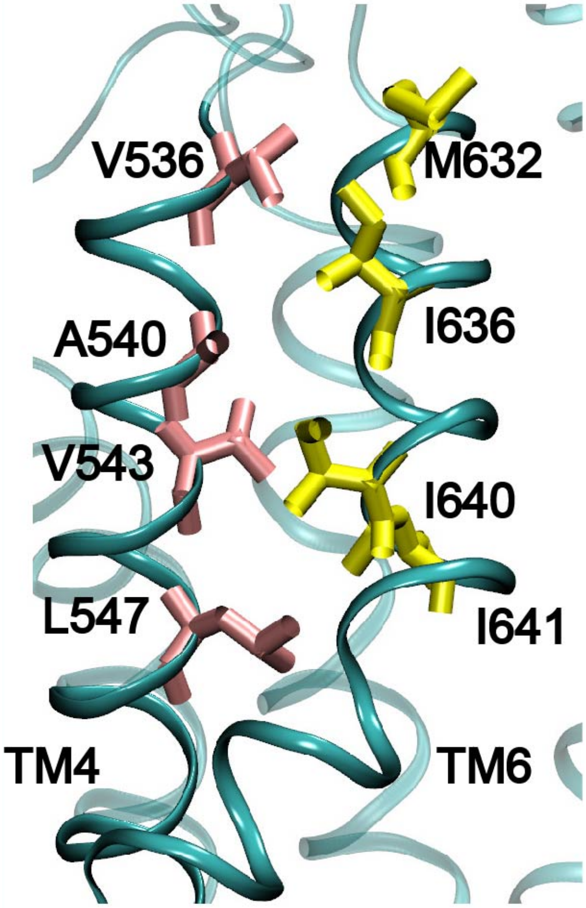


**Fig. S8** A representative snapshot of the inactive pore in the $Ca^{2+}$-free state of TMEM16A (*sim 7*, 1.0 μs). The hydrophobic residues on TMs 4 and 6 are represented as pink and yellow sticks, respectively.
